# PAX6 protein in neuromasts of the lateral line system of salamanders (*Eurycea*)

**DOI:** 10.1101/2023.10.08.561443

**Authors:** Brittany A. Dobbins, Ruben U. Tovar, Braden J. Oddo, Thomas J. Devitt, David M. Hillis, Dana M. García

## Abstract

PAX6 is well known as a transcription factor that drives eye development in animals as widely divergent as flies and mammals. In addition to its localization in eyes, PAX6 expression has been reported in the central nervous system, the pancreas, testes, Merkel cells, nasal epithelium, and developing cells of the inner ear. Here we show that PAX6 is also found in the mechanosensory neuromasts of the lateral line system in paedomorphic salamanders of the genus *Eurycea*. Using immunohistochemistry and confocal microscopy, we found that PAX6 is extranuclear, and antibody labeling is most intense in the apical appendages of the hair cells of the neuromast. This extranuclear localization raises the possibility of an as yet undescribed function for PAX6 as a cytoskeleton-associated protein.

## 1. Introduction

PAX6 is well known as a transcription factor that drives eye development in animals as widely divergent as flies and mammals. In addition to its localization in eyes, PAX6 expression has been reported in the central nervous system, pancreas, and testes (see [1]), Merkel cells in whisker follicles [2], developing nasal epithelium [3], limbal epithelial niche cells [4], and developing cells of the inner ear [5]. In these tissues, PAX6 functions as a transcriptional regulator of cell proliferation and differentiation. When its expression is suppressed, as observed in Mexican cavefish (*Astyanax mexicanus*), eye development fails to progress: apoptosis leads to involution and the disappearance of the lens and an underdeveloped retina (reviewed in [6]).

Like the genus *Astyanax*, the salamander genus *Eurycea* includes aquatic species that live at the surface, species that live underground, and species that have populations in both environments (see [7], for a recent phylogeny). For example, the San Marcos salamander (*E. nana*) lives in the surface waters emanating from the San Marcos Springs about 50 km south of Austin. The Texas blind salamander (*E. rathbuni*) inhabits the Edwards Aquifer in central Texas which periodically spews individuals to the surface via artesian wells in and around the city of San Marcos. The Cascade Caverns salamander (*E. latitans*) has both a surface and subterranean phenotype, the latter showing greatly reduced eyes along with several other characteristics typical of cave organisms.

As part of our work investigating *Pax6* expression during eye development in the surface-dwelling San Marcos salamander and the subterranean Texas blind salamander, we serendipitously observed labeling for PAX6 in the skin of both species. Here we present evidence that PAX6 is present in the mechanosensory neuromasts of the lateral line with particularly intense labeling in the apical appendages of the hair cells of the neuromast. This finding is surprising both because it is a rare instance of PAX6 labeling in the peripheral nervous system and because the labeling manifests extranuclearly, raising the possibility of an as yet undescribed function for PAX6.

## 2. Materials and methods

### 2.1 Animals

The San Marcos Aquatic Resources Center (SMARC) is operated by the United States Fish and Wildlife Service and serves as a refugium for endangered species and maintains captive, breeding populations of San Marcos salamanders (*E. nana*), Texas blind salamanders (*E. rathbuni*), as well as other threatened and endangered species (see https://www.fws.gov/office/san-marcos-aquatic-resources-center). Salamander larvae of both species (1 month post-oviposition; Fig. 1) were graciously provided by SMARC after euthanasia and fixation on site in 4% formaldehyde (derived by alkaline depolymerization of paraformaldehyde) prepared in 0.1 M phosphate buffered saline (PBS), pH 7.4.

**Fig 1.**
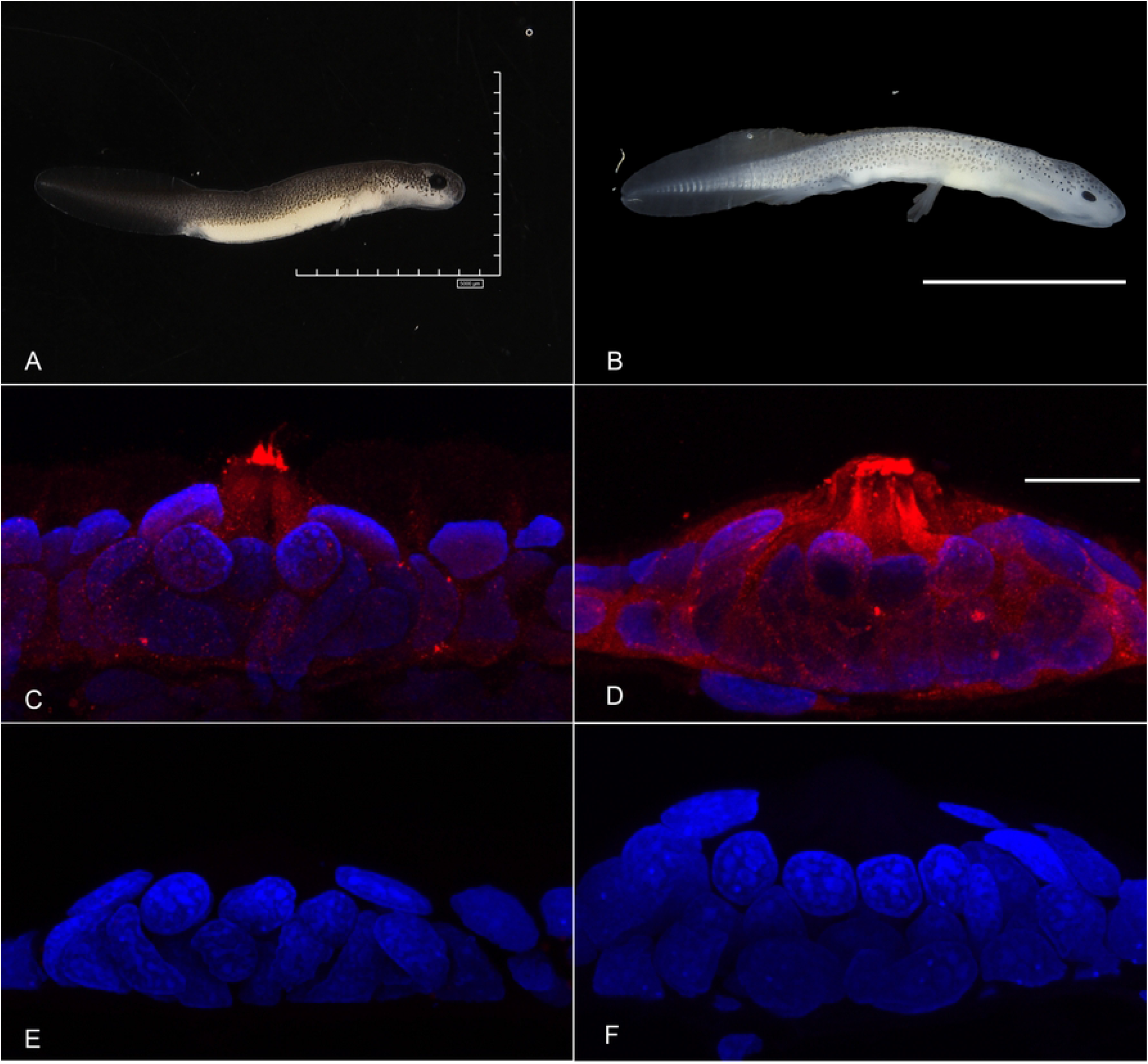
PAX6 (red) localizes to the apical appendages of hair cells of the neuromasts of the anterior lateral line system of both San Marcos salamander larvae (A, C, E) and Texas blind salamander larvae (B, D, F). Along with intense labeling of the apical appendages, diffuse labeling in the cytoplasm is also observed (C, D) along with some nuclear labeling in the San Marcos salamander’s neuromast (C). Negative controls images (no exposure to anti-PAX6 antibody) for San Marcos (E) and Texas blind (F) salamanders are presented for comparison. The neuromasts imaged are from the larvae shown in A and B. The scale bars in A and B are 5000 μm; the scale bar in D is 20 μm. Confocal images represent Z-projections acquired at the same magnification and settings. Nuclei are labeled blue with DAPI.

### 2.2 Tissue preparation and sectioning

Twenty-four hours after specimens were fixed, they were washed three times in PBS for 15 minutes/wash and then incubated at 4°C in 30% sucrose prepared in PBS with 0.1% sodium azide as an antimicrobial agent. Once specimens were cryoprotected as indicated by sinking to the bottom of the tube, images of each individual were acquired using a Hirox digital microscope prior to processing for sectioning, and specimens were assigned unique identifying numbers and stored individually at 4°C until use. Specimens were embedded in TissueTek^®^ and cut into 15 μm sections at -22°C using a Leica Cryostat equipped with a disposable blade.

Sections were collected on gelatin-coated glass slides and stored at -20°C until use.

### 2.3 Immunolabeling

Immunolabeling was accomplished as previously described [8]. In brief, sections were incubated for one hour at room temperature in PBS containing 0.05% Tween 20 (Promega, H5152) and 5% non-fat dry milk as a blocking agent. Following blocking, sections were washed three times for 10 minutes/wash with PBS containing 0.05% Tween. Sections were then incubated with anti-PAX6 antibody (Thermo Fisher/Invitrogen PA1-801, see S1 Fig. for Western Blot analysis of antibody) diluted 1:75 in PBST containing 0.1% non-fat dry milk. Slides to serve as negative controls were not incubated with anti-PAX6, but with PBST containing 0.1% non-fat dry milk. Following a two-hour incubation at room temperature, slides were washed three times for 10 minutes/wash in PBS. Sections were then incubated for one hour at room temperature with goat anti-rabbit secondary antibody conjugated to Cy5 (Thermo Fisher Scientific, A10523), diluted 1:500 in PBST containing 0.1% non-fat dry milk. Following this step, sections were washed three times for 10 minutes/wash in PBS. Coverslips were mounted onto the slides with Everbrite^TM^ Fluorescence Antifade Mounting Medium (Quartzy/Biotium, 23002) containing DAPI as a nuclear stain. Slides were stored frozen at -20°C until used.

### 2.4 Image acquisition and preparation

Images were obtained using an Olympus FV1000 Laser Scanning Confocal Microscope. Three neuromasts from each of three individuals of each of the two species were imaged. Confocal laser settings were optimized for the DAPI channel for each section. The laser settings for the Cy5 channel were optimized using a sample generated from a Texas blind salamander, and the same Cy5-settings were used for all images acquired. Images were captured as Z-stacks comprising 30 optical sections with a step size of 0.41 μm and were prepared for publication using Adobe Photoshop 24.5.0.

### 2.5 Sequence analysis

Using a comprehensive list of human PAX protein sequences and NCBI accession numbers provided by Thompson et al. [1] as a starting point, sequences of all PAX proteins were downloaded and aligned using Geneious Prime 2023.1.2. Species that shared taxonomic similarities with *Eurycea* salamanders which had PAX6 sequences were obtained as well and aligned against the human PAX6 sequences. The PAX6 sequence for axolotl (*Ambystoma mexicanus*) was acquired from a BLAST search of the axolotl proteome. We acquired the Iberian ribbed newt (*Pleurodeles walti*) sequence by BLAST using the immunogenic sequence and observing the top 100 results that exhibited the immunogenic sequence. Obtaining the Japanese fire-bellied newt (*Cynops pyrrhogaster*) PAX6 sequence was also uncovered using the BLAST with the results were filtered by the superfamily Salamandroidea and having a percent identity of 95%-100%. These sequences were aligned and figures were generated using Geneious.

## 3. Results

Immunolabeling with anti-PAX6 antibodies revealed labeling of the cells of the neuromast with particularly intense labeling of the apical appendages, in some cases lending the hair cells of the neuromast the appearance of an erupting volcano (Fig. 1). This pattern of labeling was evident in six of the nine neuromasts observed in the Texas blind salamander, and in all of the neuromasts observed in the San Marcos salamander. Labeling was also noted in the surrounding epithelial cells, especially in the samples derived from the Texas blind salamander larvae. Labeling within the nucleus was variable. In some of the negative control images, cells beneath the skin were sometimes observed to be dimly red in the Cy5 channel; however, we never observed non-specific labeling in the neuromasts (see Fig 1e, f for examples).

To address the question of whether the anti-PAX6 antibody (raised against an immunogenic sequence corresponding to mouse PAX6 and identical to human PAX6) would recognize all the isoforms of PAX6, we analyzed the sequences of all human PAX6 proteins. This analysis revealed that all fifteen isoforms of human PAX6 contain the immunogenic sequence published by ThermoFisher, specifically REEKLRNQRRQASNTPSHI (Fig 2A). Additionally, none of the other human PAX proteins contained that sequence. A search of the NCBI database using BLAST and the immunogenic sequence as the query sequence yielded 100 sequences containing the identical sequence, 95 of which were identified explicitly as PAX6 sequences. Those sequences that were not explicitly labeled as PAX6 sequences were hypothetical proteins inferred from genome projects and containing PAX6-like sequences, including from the Iberian ribbed newt (*Pleurodeles walti*) and the Japanese fire-bellied newt (*Cynops pyrrhogaster*), both species in the same superfamily as the San Marcos and Texas blind salamanders, namely Salamandroidea [9]. PAX6 sequences for axolotl (*Ambystoma mexicanus*), another member of the Salamandroidea [9], also revealed 100% identity with the immunogenic sequence.

**Fig 2.**
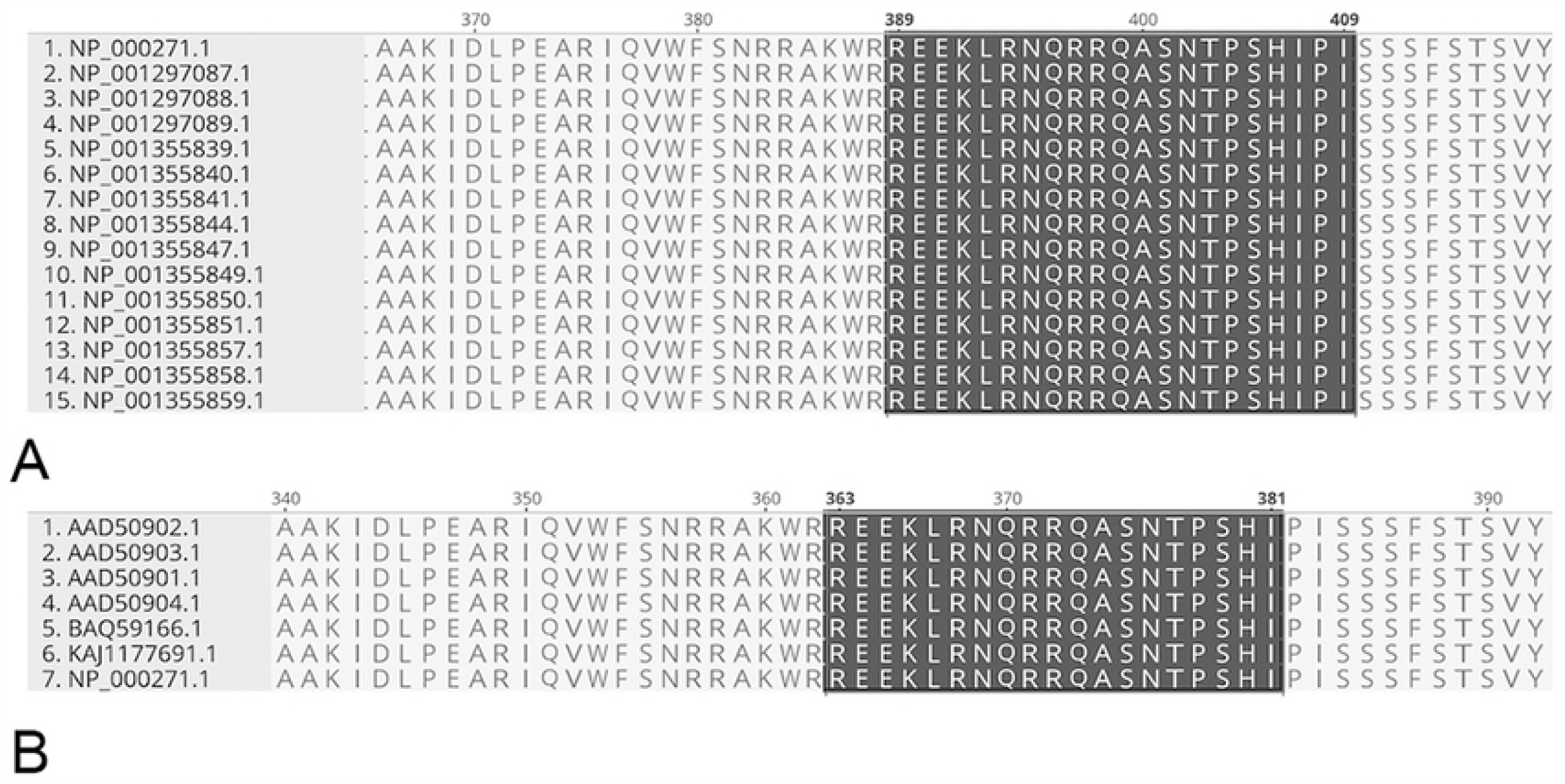
Sequence analysis supports the conclusion that anti-PAX6 antibody recognizes PAX6 protein and does not recognize other PAX proteins. (A) An alignment of human PAX6 sequences reveals that all fifteen isoforms share the immunogenic sequence (highlighted in dark grey). NCBI sequence numbers are to the left of the sequences and are arranged alphabetically by isoform. (B) The four isoforms of axolotl PAX6 (sequences 1-4) contain the immunogenic sequence as does PAX6 from the Japanese fire-bellied newt (sequence 5) and the hypothetical protein from the Iberian ribbed newt (sequence 6). Sequence 7 is human PAX6.

Comparing these sequences showed an identical region of 19 residues making up the immunogenic sequence (Fig 2B), and 92.4% identity overall by pairwise comparison. Additional differences include variability in the length of the predicted sequences; for example, in the Iberian ribbed newt, the predicted sequence has an additional 81 residues at the N-terminus. Consistent with our sequence analysis, western analysis indicated that the antibody recognized a protein that co-migrated with a protein derived from mouse retina (see S1 Fig.)

## 4. Discussion

We explored the localization of PAX6 in larvae of two species of salamanders, namely the San Marcos salamander and the Texas blind salamander. We found expression in the eye as expected (reported elsewhere) and unexpectedly in the neuromasts of the lateral line system. PAX6 localization in the nervous system has been well studied with a particular focus on the central nervous system where its nuclear localization befits its role as a transcription factor.

Parisi and Collinson [2] observed PAX6 in Merkel cells, mechanoreceptors associated with whisker follicles, and they reported its localization in developing mice as young as E16.5. Like us, they were surprised to see PAX6 localized to the cytoplasm of Merkel cells of E18.5 mice, but noted that at post-natal day 4, localization appeared to be shifting toward the nucleus.

While they did not show results from examining skin from older individuals, previous work from their lab showed cytoplasmic localization of Pax6 associated with oxidative-stress of corneal cells from adult *Pax6*^*+/-*^ mice [10]. Similarly, Parisi and Collinson [2] found that in cultured Merkel cells, they could both simulate the transition of PAX6 labeling from cytoplasmic to nuclear over time and induce nucleocytoplasmic shuttling by exposing the cells to H2O2 to induce oxidative stress [2]. The PAX6 labeling that we observed is evident in neuromasts of adult Texas blind salamanders (S2 Fig.) as well as larval. Interestingly, these salamanders share a paedomorphic life history [11], maintaining larval features throughout their lives. At the tissue level, paedomorphy is thought to occur by slowing the developmental rate of somatic tissue while maintaining gonadal tissue development and eventual maturation, a condition known as neoteny [12].

Life history may play a role in the progressive loss or gain of neuromasts. Loss of lateral line neuromast is observed in some terrestrial salamander species that undergo direct development, i.e., omitting the larval stage. During development of the lateral line, the afferent lateral line neurons innervate the skin prior to neuromast formation [13]. Interestingly, the underlying afferent lateral line neurons can still be observed within some species of dusky salamanders (genus *Desmognathus*), for example, seepage salamanders (*D. aeneus*) and pygmy salamanders (*D. wrighti*) in the absence of neuromasts [14]. The retention of afferent lateral line neurons has been interpreted as neuro-anatomical paedomorphy [15,16]. It would be interesting to explore whether neuromasts of metamorphosing salamanders that maintain a lateral line similarly evince PAX6 in their neuromasts.

Like Merkel cells [17], hair cells of the neuromasts are ectodermally derived mechanoreceptors equipped to synapse on afferent neurons of the somatosensory system (see [18] for review). The hair cells are so called because the stereocilia and kinocilium that extend from their apical surface supported by actin filaments and microtubules, respectively, to form the hair bundle (see [19]). The labeling we observed in the neuromasts was most intense in the apical appendages, which we presume to be the stereocilia because of their abundance. Studies are underway to determine whether PAX6 associates with specific cytoskeletal elements in these structures.

## 5. Conclusions

We have shown here that anti-PAX6 antibodies label the neuromasts of the San Marcos salamander and the Texas blind salamander, localizing in both the nuclei and the apical appendages of the hair cells. Sequence analysis supports our inference that the antibody is recognizing PAX6 and not a similar protein. Future studies will be directed toward better resolving the precise association of PAX6 protein with structures in the hair cells.

## Acknowledgements

We are grateful to the San Marcos Aquatic Resources Center (United States Fish and Wildlife Service) for donation of salamander larvae. The staff at the Applied Research Service Center (ARSC) of Texas State University, especially Mrs. Alissa Savage, Mr. Jacob Bisbal and Mr. Jacob Armitage, made every accommodation possible to assure BAD was trained efficiently on the confocal and digital microscopes during her tenure as an NSF Summer Research Fellow.

Mrs. Savage continues to provide helpful, technical assistance.

## Supporting information

**S1 Fig. Western Blot Analysis**. Tissue lysates were obtained from embryos of *E. rathbuni* and *E. nana* as well as adult mouse eye to serve as a positive control. A western blot analysis using our primary PAX6 antibody suggests that the antibody labels a protein of the expected molecular weight of PAX6 in all three taxa.

**S2 Fig. PAX6 protein localizes to the apical appendages of hair cells in neuromasts of adult Texas blind salamanders.** Anti-PAX6 antibodies label apical appendages (arrowhead) of hair cells in the neuromast of an adult Texas blind salamander. The scale bar in the image is 20 μm, and the image represents a Z-projection acquired at the same magnification and settings as used in Fig 1. Nuclei are labeled blue with DAPI.

